# Accelerated learning of a noninvasive human brain-computer interface via manifold geometry

**DOI:** 10.1101/2025.03.29.646109

**Authors:** Erica L. Busch, E. Chandra Fincke, Guillaume Lajoie, Smita Krishnaswamy, Nicholas B. Turk-Browne

## Abstract

Brain-computer interfaces (BCIs) promise to restore and enhance a wide range of human capabilities. However, a barrier to the adoption of BCIs is how long it can take users to learn to control them. We hypothesized that human BCI learning could be accelerated by leveraging the naturally occurring geometric structure of brain activity, or its intrinsic manifold, extracted using a data-diffusion process. We trained participants on a noninvasive BCI that allowed them to gain real-time control of an avatar in a virtual reality game by modulating functional magnetic resonance imaging (fMRI) activity in brain regions that support spatial navigation. We then perturbed the mapping between fMRI activity patterns and the movement of the avatar to test our manifold hypothesis. When the new mapping respected the intrinsic manifold, participants succeeded in regaining control of the BCI by aligning their brain activity within the manifold. When the new mapping sdid not respect the intrinsic manifold, participants could not learn to control the avatar again. These findings show that the manifold geometry of brain activity constrains human learning of a complex cognitive task in higher-order brain regions. Manifold geometry may be a critical ingredient for unlocking the potential of future human neurotechnologies.

## Introduction

Brain-computer interfaces (BCIs) allow users to control external devices like computer displays, communication tools, or robotic effectors with their brain activity [1–8]. Progress in developing BCIs for neural rehabilitation and cognitive augmentation in humans has been hindered by challenges with neural decoding and user training [8–11]. As such, current human BCIs are prohibitively slow for users to learn and require frequent calibration to maintain performance, and even still many “non-responders” are never able to gain control [12–18].

In this project, we design and validate a computational framework that enhances human BCI learning. We hypothesized that some brain states are easier for people to generate, and that tailoring training to these brain states will facilitate BCI learning. To test this hypothesis, we trained healthy human participants to control a video game with their brain activity measured noninvasively with closed-loop, real-time functional magnetic resonance imaging (rt-fMRI). This technique allowed us to capture high-resolution, whole-brain activity, analyze the data, and update the game display based on the results every 2 s (the rate of fMRI data acquisition). rt-fMRI has been used in the past as a form of noninvasive BCI for neurofeedback [19–23], yet suffers the same challenges as other BCIs of slow learning or non-response [18, 24]. Within this system, we perturbed the relationship between measured brain activity and the video game display to probe how humans learn to generate different brain states.

Recent studies on neural prosthetics in nonhuman primates provide support for this approach. In these studies, monkeys were trained to control a cursor based on invasive multi-unit recordings from primary motor cortex. Learning of this task occurred more rapidly when the target brain states conformed to the naturally occurring geometry, or “intrinsic manifold”, of the neural population activity [25–28]. This kind of manifold-based BCI learning also promotes more efficient transfer between similar brain states and supports more stable performance over time [29–35]. Together, these foundational findings suggest the importance of extracting the intrinsic manifold of brain regions to determine which states to train in human BCI learning.

Our framework represents an advance over prior work in several ways. First, given the integral role of higher-order cognitive processes in successful BCI learning [16, 36], we targeted a network of brain regions linked to spatial navigation and goal-directed behavior rather than sensorimotor regions. Second, to engage these processes during BCI learning, we designed a novel task in which participants navigated an avatar through a virtual reality (VR) arena (Figures 1, S1). This focus on modulating higher-order brain regions to control complex behavior lends insight to future applications of BCIs for enriching human cognition. Third, given the high dimensionality and noise inherent to fMRI and other noninvasive techniques, we developed state-of-the-art algorithms for learning and extending the intrinsic manifold. Data-diffusion methods have found success in applied mathematics for discovering nonlinear structure in complex biomedical processes [37–39]. We optimized a variant for fMRI — temporal potential of heat diffusion for affinity-based transition embedding (T-PHATE) — that learns a lower dimensional manifold for each participant and highlights brain states related to cognition and behavior better than other dimensionality reduction approaches [40], including the linear methods used in most prior BCI studies.

**Figure 1:**
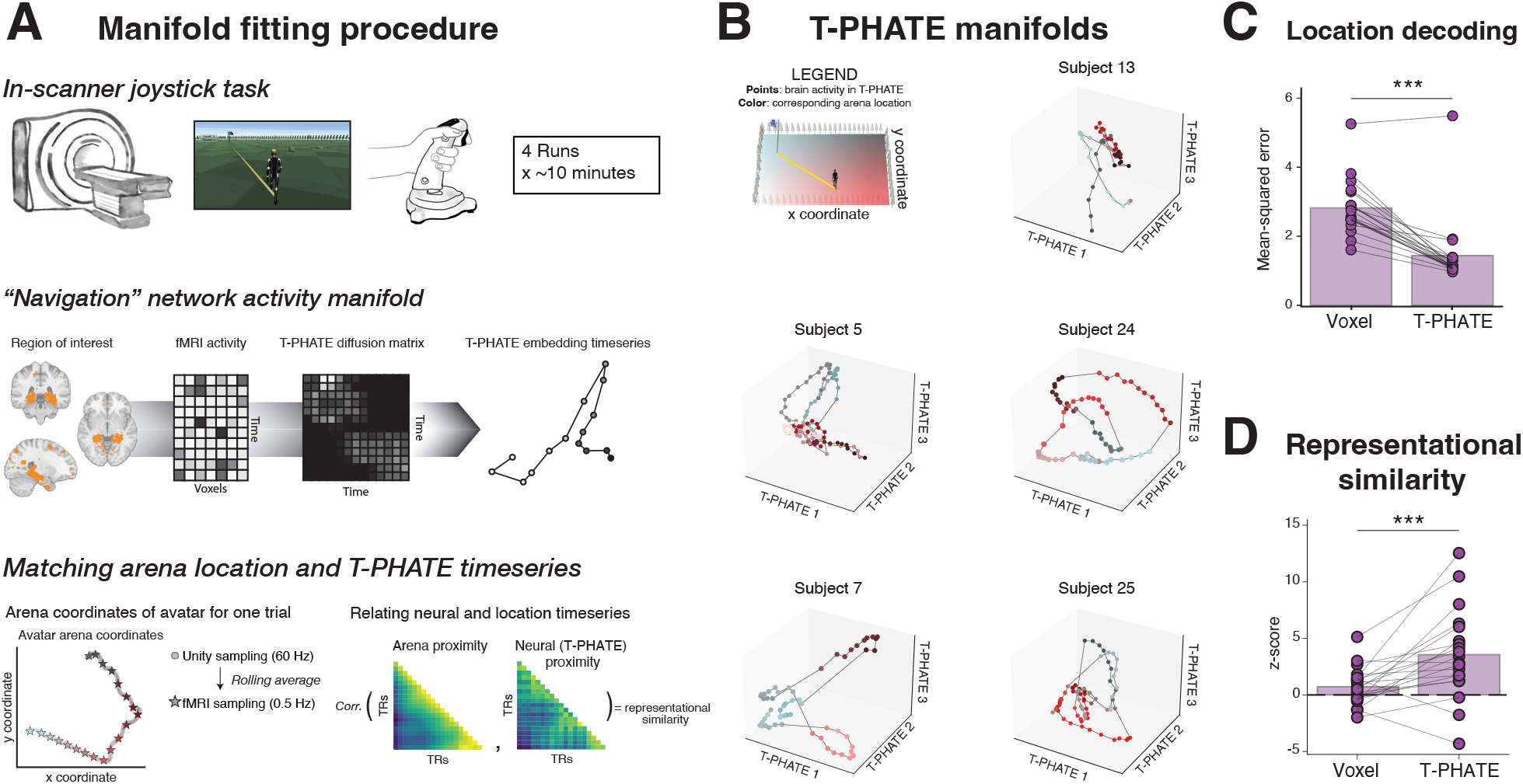
Manifold learning and validation. (A) Participants practiced self-guided navigation by steering an avatar with a joystick during a video game task (Unity3d) while undergoing fMRI scanning. Multi-voxel fMRI patterns were extracted during this task from a network of brain regions implicated in spatial navigation. These patterns were then embedded into a low-dimensional manifold using the T-PHATE algorithm. Each data point is a single fMRI sample visualized in 3 T-PHATE dimensions, with consecutive timepoints connected. For the main analyses, data were embedded into 20 T-PHATE dimensions. The avatar’s arena location was downsampled to 0.5 Hz using average pooling to match the temporal resolution of fMRI acquisition and shifted 2 samples (4 s) to correct for the hemodynamic lag. (B) T-PHATE embeddings of fMRI data reflect the structure of the video game arena. Timeseries for five participants during four consecutive trials of the joystick task are depicted, with timepoints colored by the coordinate location of the avatar in the game arena. (C) Cross-validated regression models were used to decode the avatar’s location from brain data (*N* = 20). These predictions were more accurate when data were embedded in the T-PHATE manifold relative to the original voxel space. (D) Representational similarity was used to relate distances between brain activity patterns to distances between arena locations (*N* = 20). The representational geometry of T-PHATE embeddings better matched the location geometry than the original voxel space. *** *P <* 0.001.

Participants in our study completed four fMRI sessions. The first was used to learn the participant’s intrinsic manifold with T-PHATE while they practiced navigating the avatar with a joystick. The remaining three sessions tested how manifold geometry constrained the participant’s ability to control the avatar by modulating distributed activity patterns in navigation-related brain regions. Each session required the participant to learn a new BCI mapping between fMRI activity and the avatar’s heading direction. The second session used an “intuitive mapping” (IM), which best captured the participant’s intrinsic manifold. The third and fourth sessions (order counterbalanced) critically tested perturbations of the mapping that were within or outside the intrinsic manifold (Figure 2). As hypothesized, participants rapidly learned the IM and the “within-manifold perturbation” (WMP), which respected the geometry of their intrinsic manifold, but failed to learn the “outside-manifold perturbation” (OMP), which did not respect the manifold. Successful BCI learning led to a realignment of brain activity along the manifold and to enhanced decoding of task information.

**Figure 2:**
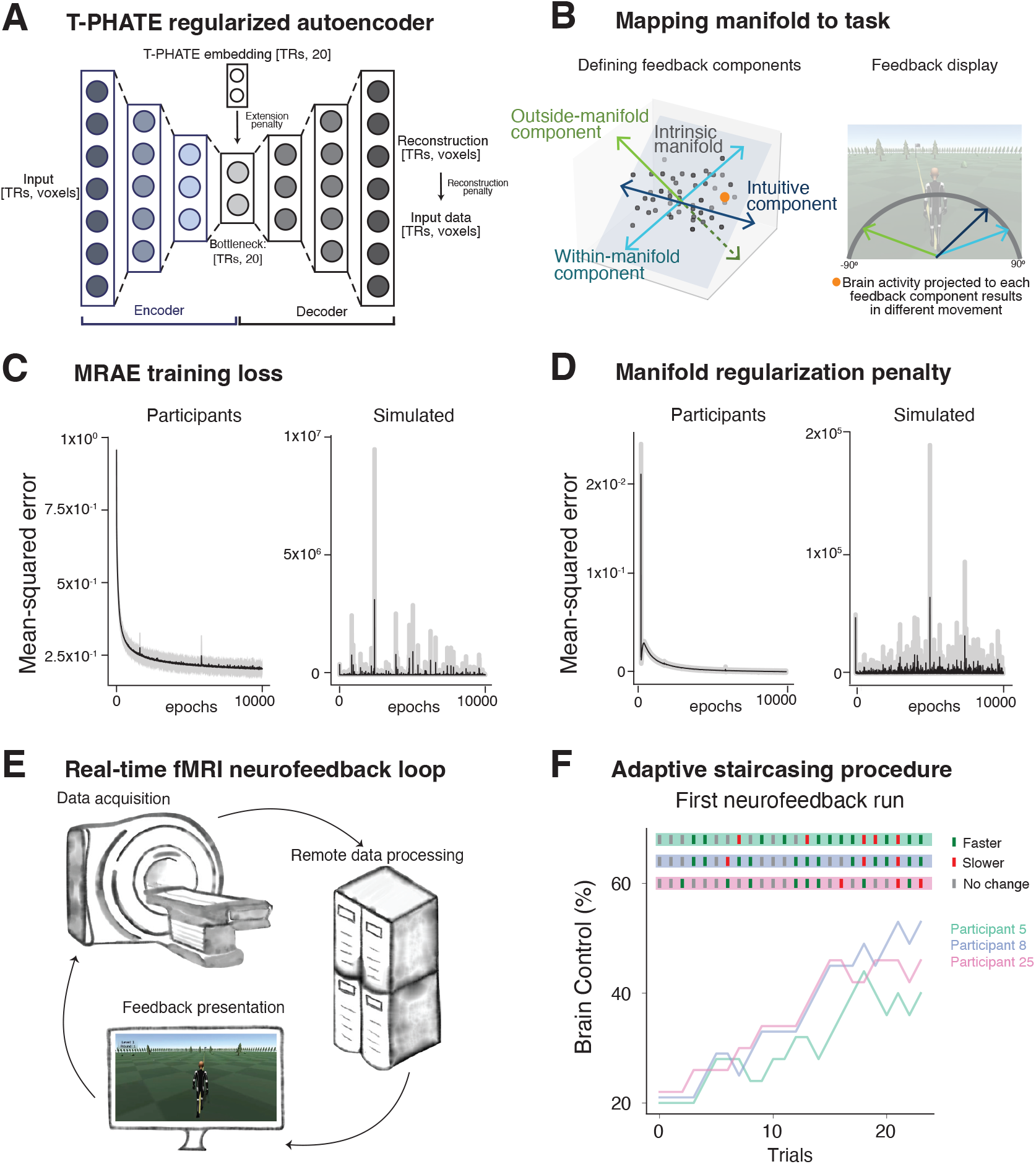
Real-time manifold extension and neurofeedback. (A) A manifold-regularized autoencoder (MRAE) uses a T-PHATE layer to penalize the latent space of the autoencoder to reflect the input data’s T-PHATE manifold geometry. (B) The intrinsic manifold (gray plane) of a dataset (black points) with an “intuitive” component (capturing a main dimension of variance on the manifold), a “within-manifold” component (a different, highly explanatory component on the manifold), and an “outside-manifold” component (explaining minimal variance). The same data point (orange) projected onto each component (left) maps to three distinct angles of movement in the task (right). (C) The training loss for MRAE combines a reconstruction penalty plus a manifold regularization penalty, shown for real participants (left) and simulated fMRI data as a null comparison (right). (D) Plotting the manifold regularization penalty component of the training loss separately. Loss functions are visualized as the mean (line) and 95% confidence interval (band) of the mean across participants. (E) Closed-loop procedure used *rt-cloud* [43] software to transfer, analyze, and administer continuous feedback based on fMRI scans acquired every 2 s. (F) The proportion of control that a participant’s brain activity exerts over the avatar’s movement — the *Brain Control* parameter — is adjusted at the end of each trial with an adaptive staircasing procedure. *Brain Control* increases with improved accuracy (green dashes) and decreases with worsening accuracy (red dashes). The accuracies and staircasing curves of three representative participants are depicted for the first neurofeedback run.

## Results

### Discovering the manifold of human brain regions

Our initial goal was to learn a meaningful manifold of brain activity that could be used in later sessions to guide BCI learning. We first used the T-PHATE approach [40] to validate that activity measured from a network of navigation-related brain regions contained dynamic information about the joystick task. A key use of manifold learning methods is for visual exploration of high-dimensional data in lower dimensions [39], so we embedded each participant’s fMRI timeseries data for all trials of the joystick task into three dimensions using T-PHATE and labeled each timepoint by the avatar’s corresponding coordinate in the game arena (Figure 1A). Showing that the brain activity embedded with T-PHATE reflected the structure of the arena, points were clustered according to their proximity in the arena and trajectories through the points over time traversed the arena smoothly (Figure 1B).

Prior work has shown that T-PHATE embeddings do not just aid in visualization but also improve extraction of task-related information [40, 41]. We validated that the network of brain regions selected for neurofeedback represented navigation-related information during the joystick task, and that this information was represented more clearly after the data were embedded with T-PHATE. For each participant, we trained cross-validated linear regression models on the voxel-resolution and T-PHATE-embedded fMRI timeseries data (here, 20-D embeddings) to predict the avatar’s arena coordinates at each timepoint. Regression models were scored as the mean-squared error between model-predicted and true coordinates in held-out data, such that lower error indicated superior decoding. Models trained on T-PHATE embeddings were more accurate than those trained on voxel-resolution data (*P* = 0.0002), supporting that the representation afforded by manifold learning enhanced access to task-related information (Figure 1C).

Beyond predicting individual locations, we tested whether the representational geometry of brain activity in neurofeedback regions reflected arena geometry. In other words, were proximal coordinates represented more similarly by the brain than more distant ones, and does embedding with T-PHATE clarify this representational structure? We used a Mantel test to evaluate representational similarity as z-scores relative to a null distribution [42] and found that distance in the game arena was more correlated with pattern similarity of T-PHATE embeddings than of voxel-resolution activity (*P* = 0.0008; Figure 1D).

### Extending the manifold to new samples for neurofeedback

Despite the benefits of manifold learning for modeling low-dimensional neural dynamics, a key drawback of most nonlinear dimensionality reduction methods is that the learned dimensions cannot be easily extended to untrained data [39]. To circumvent this limitation, we developed a manifold-regularized autoencoder (MRAE) that could embed a participant’s untrained data in their respective T-PHATE manifold in real-time [44, 45]. MRAE is trained to optimize for two measures of model fit: (1) a reconstruction error penalty, which uses mean-squared error (MSE) to quantify how well the autoencoder reproduces the original data; and (2) a manifold regularization penalty, which quantifies the error between where the encoder places a sample and its true location on the manifold. These penalties are combined into a single loss function to train MRAE. This training loss decreased rapidly and plateaued by 6,000 training epochs (mean across participants: *MSE* = 0.21, *s*.*e*.*m* = 0.0093; Figure 2C, left). The manifold regularization penalty on its own converged by 5,000 training epochs (*MSE* = 0.0015, *s*.*e*.*m*. = 0.00004; Figure 2D, left).

To establish a null baseline, we conducted the full multi-session experiment, including the neurofeedback pipeline calibration and real-time procedure, on fake participants whose simulated fMRI activity consisted only of realistic noise estimated from the brain activity of neurofeedback participants [46]. The data did not contain task-related responses during the calibration session or learning effects during the neurofeedback sessions, and thus we did not expect T-PHATE to learn a useful manifold or for the MRAE training to succeed. Indeed, the overall MRAE training loss (*MSE >* 1, 500, *s*.*e*.*m. >* 500; Figure 2C, right) and the subscore for the manifold regularization penalty (*MSE >* 600, *s*.*e*.*m. >* 40; Figure 2D, right) neither decreased nor converged. Thus, the neural manifolds of real participants engaged in the joystick task reflected learnable, extensible structure, which was not found in simulated brains devoid of task-driven signal (Supplemental Figures S2, S3).

### Manifold-constrained learning of BCI control

We quantified BCI learning by the increase in *Brain Control*, a confidence parameter that tracked the ability of participants to direct the avatar toward the goal efficiently with their brain activity (Figure 2F). Specifically, *Brain Control* is the staircased proportion [47] of the avatar’s movement direction dictated by the angle decoded from the brain; the remaining proportion reflects the angle straight toward the goal, providing assistive guidance in the right direction. Change in *Brain Control* (Δ*BrainControl*) was computed at each trial relative to the first trial of the session (Figure 3A). In the first neurofeedback session, BCI learning of the intuitive mapping (IM) occurred rapidly, reaching significance after trial 4 and plateaued around the final 15 trials. Adjusting to the within-manifold perturbation (WMP) in the second or third neurofeedback session (counterbalanced) led to a slower but steady increase in *Brain Control*, reaching significance around trial 30. Participants did not adjust to the outside-manifold perturbation (OMP), as indicated by the lack of change in *Brain Control* from baseline.

**Figure 3:**
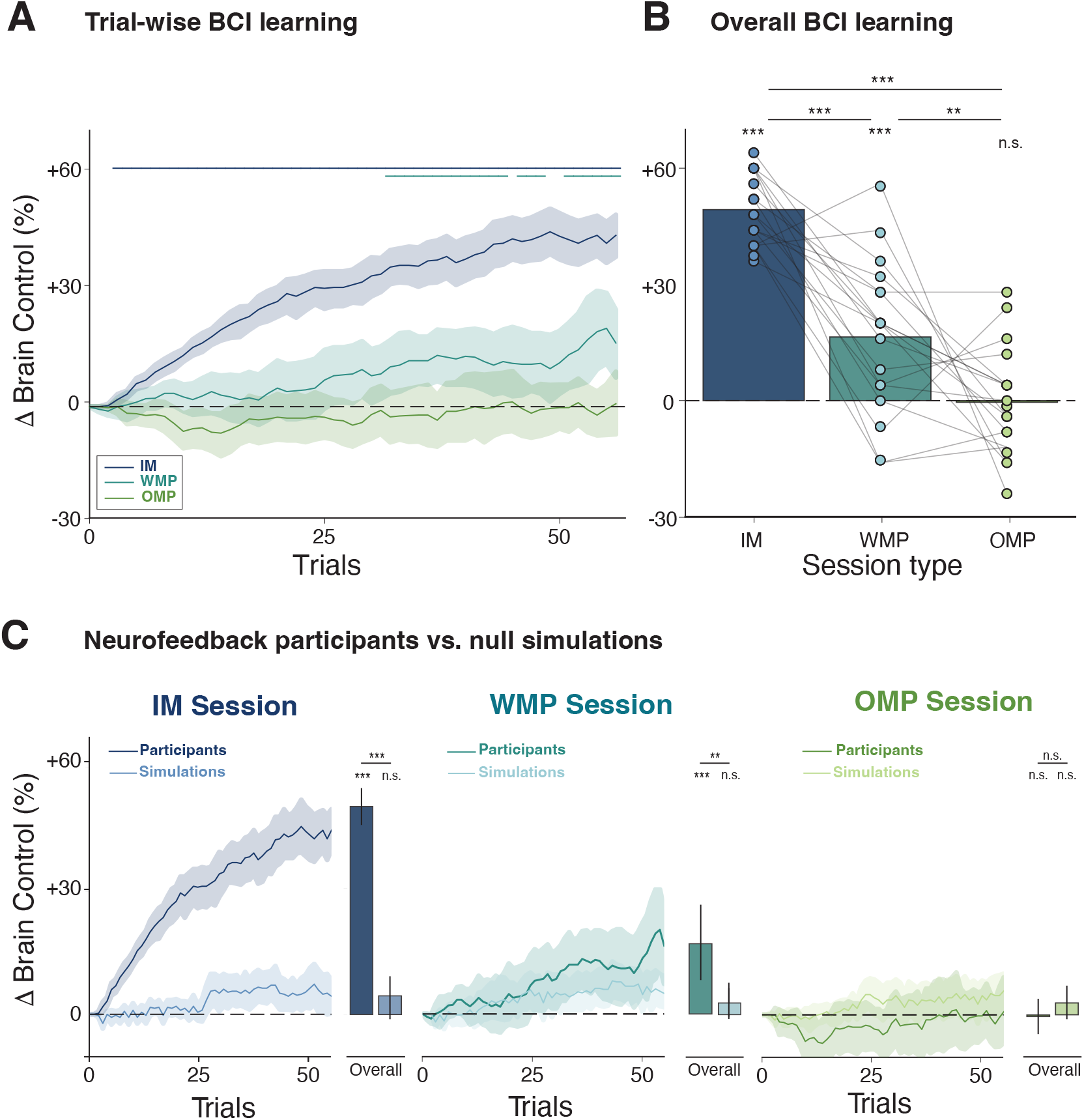
BCI learning of different manifold components. (A) Learning curves of the change in *Brain Control* across training trials for each session type. Lines show mean across participants (*N* = 18) and bands show the 95% confidence interval (CI) of the mean. The number of trials (55) depicted on the x-axis corresponds to the minimal trial count successfully completed by all participants, across all session types (participants who performed better could complete a few additional trials in the same amount of time, not shown here so the data reflect the full sample). Horizontal lines at the top indicate where the mean was significantly greater than chance (Δ*BrainControl >* 0, *P <* 0.05) for each session type (non-parametric cluster-based correction). (B) Total change in *Brain Control* from the first trial to the last trial completed in each session type, for each participant. The total number of completed trials varied slightly by participant and session. Bars represent the mean across participants; points represent individual participants; lines connect the points from the same participant across the three sessions. (C) Comparing trial-wise and overall BCI learning by session type across real (darker; *N* = 18) and null (lighter; *N* = 20) participants. Lines show the mean across participants and bands show the 95% CI of the mean. Bars indicate mean overall learning and error bars represent 95% CI of the mean. *** *P <* 0.001, ** *P <* 0.01, * *P <* 0.05, n.s. *P ≥* 0.1.

Overall BCI learning was computed as Δ*BrainControl* from the first to the final trial of each session type (Figure 3B). There was a significant positive learning effect for the IM session (mean across participants: Δ*BrainControl* = 49.3, *s*.*e*.*m*. = 2.0, *P <* 0.0001, 95%*CI* = [45.3, 53.1]) and WMP session (Δ*BrainControl* = 16.4, *s*.*e*.*m*. = 4.5, *P* = 0.0002, 95%*CI* = [7.3, 25.3]), but not for the OMP session (Δ*BrainControl* = *−*0.4, *s*.*e*.*m*. = 3.2, *P* = 0.556, 95%*CI* = [*−*6.2, 6.4]). The increase was significantly greater in the IM session than both the WMP session (*P <* 0.001; 95%*CI* = [22.0, 42.4]) and OMP session (*P <* 0.001; 95%*CI* = [41.8, 55.6]), as well as in the WMP session compared with the OMP session (*P* = 0.002; 95%*CI* = [5.6, 30.2]).

BCI learning effects in neurofeedback participants were benchmarked against simulated null fMRI participants that underwent identical procedures without the possibility for learning (Figure 3C). As expected, these simulated data did not show a significant increase in *Brain Control* in any of the session types (*Ps >* 0.1). This analysis verifies that the observed BCI learning effects in the real participants were not an artifact of the staircasing procedure or real-time fMRI processing pipeline.

### Realignment of neural activity along manifold

We hypothesized that the mechanism for BCI learning is the alignment of brain activity with the manifold component being trained. That is, participants can gain control over the avatar by realigning their brain activity in high-dimensional space so that variance maps more precisely to changes in movement direction. To assess this neural realignment, we calculated the percentage of total variance in the fMRI signal of each run that could be explained by the feedback component for each session type (Figure 4A). We then compared the change in this percentage of explained variance (Δ*PEV*) between the first and final runs of each session.

**Figure 4:**
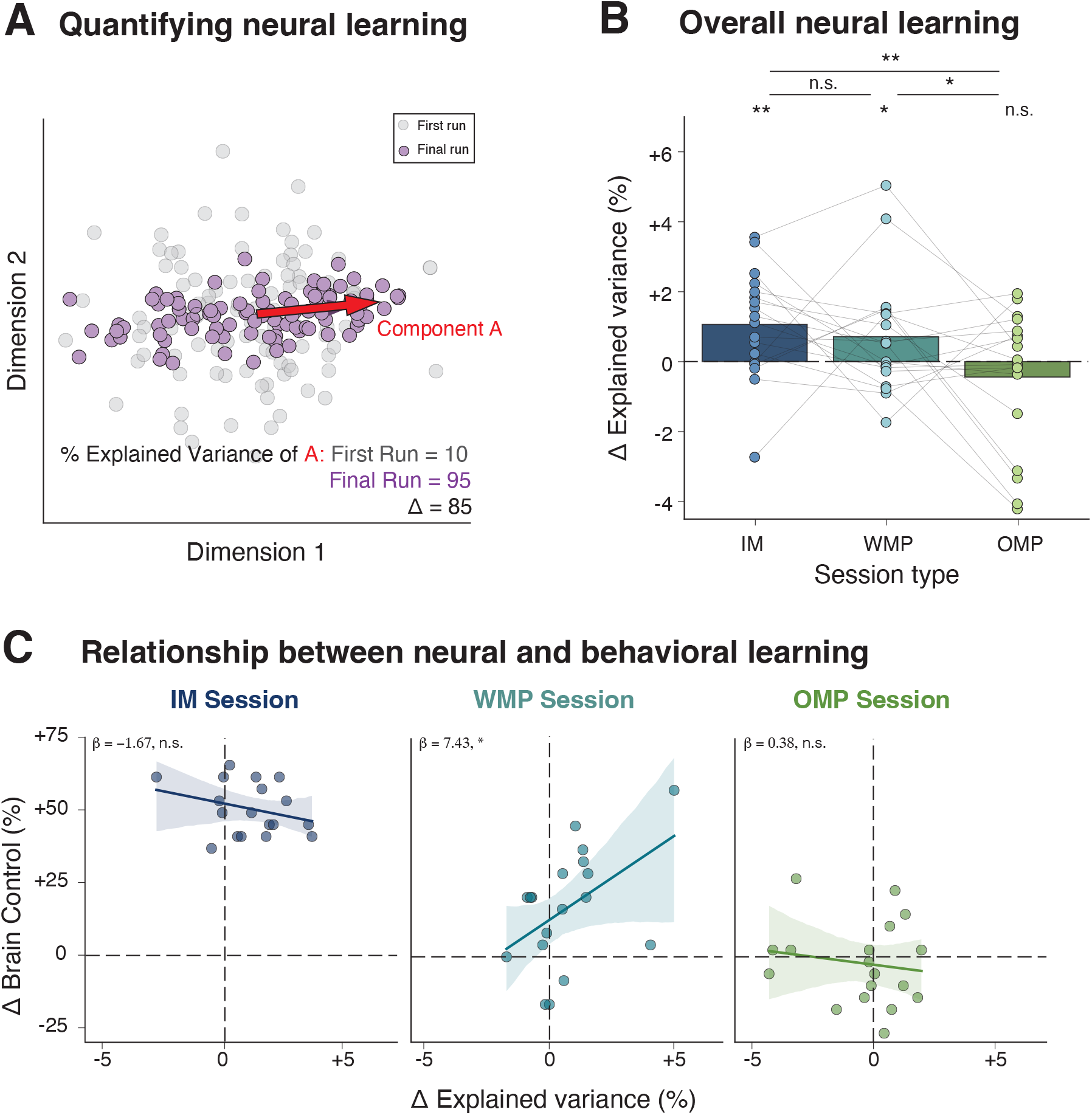
Changes in neural activity supporting BCI learning. (A) Simulated data showing a component (red vector) that explains 10% of the explainable variance initially (gray points) but 95% after learning (purple points). (B) Change in the percentage of explained variance (Δ*PEV*) over neurofeedback training for each session type. Bars represent the mean across participants; points represent individual participants; lines connect the points from the same participant across the three sessions. (C) Predicting BCI learning (Δ*BrainControl*) from changes in neural realignment (Δ*PEV*). Betas, P-values, and confidence intervals from linear mixed effects model with participant and counterbalancing order as random effects. ** *P <* 0.01, * *P <* 0.05, n.s. *P ≥* 0.1

Neural activity became more aligned with the IM component (mean across participants: Δ*PEV* = 1.1, *s*.*e*.*m*. = 0.4, *P* = 0.002, 95%*CI* = [0.3, 1.7]) and the WMP component (Δ*PEV* = 0.7, *s*.*e*.*m*. = 0.4, *P* = 0.019, 95%*CI* = [0.1, 1.6]) during their respective neurofeedback sessions, but not with the OMP component (Δ*PEV* = *−*0.4, *s*.*e*.*m*. = 0.5, *P* = 0.83, 95%*CI* = [*−*1.5, 0.3]) (Figure 4B). Importantly, these increases in alignment for IM and WMP were selective to the trained component: there was no increase in alignment with WMP or OMP components in the IM session, nor with IM or OMP components in the WMP session (Figure S4). The amount of neural realignment was comparable between IM and WMP components in their respective sessions (*P* = 0.50, 95%*CI* = [*−*0.7, 1.3]), and greater than for the OMP component in its session (vs. IM: *P* = 0.001, 95%*CI* = [0.5, 2.5]; vs. WMP: *P* = 0.021, 95%*CI* = [0.06, 2.32]).

### Relationship between BCI learning and neural realignment

Neural activity became more aligned with “on-manifold” components (IM and WMP) during neurofeedback training. We used a linear mixed effects model to predict BCI learning (Δ*BrainControl*) from this neural realignment (Δ*PEV*), while controlling for participant and counterbalancing order as random effects. There was no significant relationship between the two metrics for the IM (*β* = *−*1.67, *P* = 0.456, 95%*CI* = [*−*6.1, 2.8]) or OMP (*β* = 0.38, *P* = 0.89, 95%*CI* = [*−*5.28, 6.04]) sessions. In the WMP session, however, neural realignment reliably predicted BCI learning: *β* = 7.43, *P* = 0.017, 95%*CI* = [1.40, 13.47] (Figure 4C).

We interpret this lack of relationship for IM as reflecting the fact that the IM component was already the strongest source of variance and thus that additional neural alignment did not further benefit BCI control. Such realignment was not possible for the OMP, and there was no significant BCI learning or neural realignment for this session on average. The WMP session always occurred after the IM session (either immediately or after OMP) and so participants had to realign activity from the IM to WMP components in order to gain control. This realignment was possible, given that WMP remained on the manifold, and whether it occurred determined the amount of BCI learning in this session.

We also investigated another signature of BCI learning that was unique to the WMP condition: improved task representation. Using an exploratory whole-brain analysis to consider non-target brain regions, we found that WMP training resulted in significantly improved task decoding in primary motor cortex and regions associated executive control and cognitive strategies, including dorsolateral and ventrolateral prefrontal cortex. In contrast, there were no regions that showed significantly improved representations during the IM or OMP sessions (Supplemental Figure S5). Together, these results indicate that learning a new mapping on an existing manifold alters neural activity within the targeted regions and throughout cortex.

## Discussion

The proliferation of human BCI technologies has been hindered to date by the slowness and variability of learning across individuals. These challenges persist whether the BCI uses invasive or noninvasive neural recordings [8]. Prior real-time fMRI neurofeedback studies have typically required 4–10 training sessions to achieve changes in perception and cognition [19–23, 48, 49]. Moreover, one-third of neurofeedback users remain unable to change their brain activity [18, 24]. In the current study, all participants successfully learned to self-regulate brain activity to navigate a virtual avatar when neurofeedback was administered along their intrinsic neural manifold. Once participants achieved this control, they were able to re-learn — within one session — how to navigate the avatar after a perturbation in the mapping that stayed within their manifold. However, participants were unable to re-learn control with the same amount of training when they had to generate outside-manifold brain activity.

This accelerated on-manifold learning parallels studies in which humans and nonhuman primates learned a novel behavior (cursor control) via an invasive BCI in the motor cortex [25, 27–29]. One key difference in the current study is the use of a diffusion-based manifold learning method that was necessitated by the autocorrelation and noise of fMRI data. Given the ability of T-PHATE to recover complex signals within noisy samples (Supplemental Figure S6), it may benefit a broader range of BCI applications, including invasive studies of humans and nonhuman primates focused on language and sensorimotor brain regions, which have so far relied on simpler representations or linear methods [2, 4–6, 25–29, 50].

One reason that manifold learning methods like T-PHATE can accelerate BCI learning may be that they yield high-quality feedback that promotes durable neural plasticity. Namely, previous work found that BCI learning of within-manifold perturbations is supported by neural *reassociation* – the remapping of existing activity patterns to a new behavior [26]. In contrast, BCI learning in the current study was supported by neural *realignment* – the generation of novel activity patterns to drive a behavior. Although neural reassociation can occur more quickly, neural realignment is the behaviorally optimal solution [26]. In addition to using T-PHATE to extract a richer, more informative manifold [40], our framework may have enabled rapid neural realignment by encouraging incremental learning with a staircasing procedure, which has been linked to generating novel activity patterns [9, 27, 51–55].

The fact that participants failed to learn an outside-manifold perturbation is consistent with computational models and studies of nonhuman primates, in which such learning can take approximately 10 times as much training as a within-manifold perturbation [27, 52, 54, 56]. This suggests a potential explanation for why prior attempts at BCI learning with real-time fMRI neurofeedback have required numerous sessions to achieve success in even a majority of participants. By designating neurofeedback targets agnostic to the intrinsic manifold, these studies may have inadvertently asked participants to generate outside-manifold activity patterns. That may be desirable in some cases, for example if the goal is to change a disordered manifold to alleviate psychiatric or neurological symptoms [57, 58]; in such cases, better characterizing the initial manifold would support incremental training of outside-manifold activity. However, if the goal is to interface the human brain with computer technologies for communication or occupational applications, rather than altering the manifold itself, leveraging the intrinsic manifold will be most efficient. Our findings offer a manifold-guided route for improving basic and translational BCIs. We anticipate that these improvements will further increase the efficacy of BCIs on wearable devices that are cheaper and easier to scale for public benefit.

## Methods

### Participants

Twenty young adults were recruited from the New Haven community. All participants provided informed consent to an experimental protocol approved by the Yale University Institutional Review Board. Data from two participants were excluded from neurofeedback analyses (one for falling asleep in multiple fMRI runs and one because of a scanner malfunction). This resulted in a final cohort of 18 participants (9 female; age range = 19 *−* 35 years, mean age = 25.8 years). This sample size is greater than, or comparable with, other multi-day real-time fMRI studies [19, 20, 22, 23, 49, 59–61]. There was no separate control group because all comparisons and baselines are within-participant over time and across repeated-measures conditions. Participants completed four or five fMRI sessions lasting 1–1.5 hours each and were compensated for their participation ($20/hour plus retention incentives: $5/session cumulative bonus for each return session and $40 for completion).

### Task design

The stimulus was a custom video game programmed with C# for Unity (https://unity3d.com/; Version 2019.4.12f1) with a human avatar in a virtual outdoor arena environment. Participants were tasked with steering this avatar from its initial location to a flag across the arena. A yellow line was always visible, highlighting the shortest path from the avatars current location to the goal location. The starting location of the avatar in the arena was randomly generated at the start of each trial and the flag was placed at the same distance (but random angle) from this location for each trial within a run. The distance between the initial location and the goal increased by 5% per run. Landscape objects (shrubs, trees, and rocks) were randomly placed throughout the environment as landmarks but never obstructed the path between initial and goal locations.

The video game was presented in the scanner on a screen at the back of the bore using a rear projector (1, 920 *×* 1, 068-pixel resolution; 60-Hz refresh rate). Participants viewed the display through a mirror mounted on the MRI headcoil. The game was run through Unity on a Linux computer. The fMRI scanner triggers were collected via PsychoPy (Version 2022.2.4) running in parallel with Unity on the Linux machine. Custom Python software connected PsychoPy and Unity via a Transmission Control Protocol (TCP) port to synchronize the timing of game events with scanner triggers. A message was sent from PsychoPy to Unity to start the game when the first scanner trigger was received, and messages were sent from Unity to PsychoPy at the start and end of each trial. At the start of each neurofeedback run, task instructions were displayed to participants for 20 s.

### Data acquisition

Data were acquired using a 3T Siemens Prisma scanner with a 64-channel head coil at the Brain Imaging Center at Yale University. For joystick and neurofeedback functional runs, BOLD data were collected with an echo-planar imaging (EPI) sequence (TR = 2 s; TE = 30 ms; voxel size = 3 mm isotropic; FA = 8^*°*^; IPAT GRAPPA acceleration factor = 2; distance factor = 25%). Functional runs were of variable length due to the self-guided nature of the task, with approximately 300 volumes (10 minutes) collected per run. The experimenter aimed to end the task manually at the completion of a trial closest to 300 TRs, resulting in an average of 301.4 TRs per run (range = 150 *−* 336).

After all functional runs were complete, we acquired two spin-echo field map volumes in opposite phase encoding directions for field map correction (TR = 5 s; TE = 80 ms; otherwise matching the EPI sequences). During the joystick session, we collected two high-resolution anatomical images for spatial registration: a 3D T1-weighted magnetization-prepared rapid acquisition gradient echo (MPRAGE) scan with an IPAT GRAPPA acceleration factor of 2 (TR = 2.5 s; TE = 2.9 ms; voxel size = 1 mm isotropic; FA = 8^*°*^; 176 sagittal slices), and a 3D T2-weighted fast spin echo scan with variable flip angle and IPAT GRAPPA acceleration factor of 2 (TR = 3.2 s; TE = 565 ms; voxel size = 1 mm isotropic; 176 sagittal slices).

### Neurosynth navigation network

Neurofeedback training targeted a large network of voxels involved in self-directed navigation downloaded from https://neurosynth.org [62]. We entered the search term “navigation,” and Neurosynth synthesized activation coordinates from 77 published studies with “navigation” in the title or abstract. We downloaded the association test *z*-statistic map, which displays voxels that are reported more frequently in articles with (vs. without) this term and then performs false discovery rate (FDR) correction (*FDR* = 0.01). During preprocessing, we derived a transformation from the participant’s native space functional image to the 2-mm Montreal Neurological Institute (MNI) standard brain. We inverted this transformation to align the navigation mask from MNI space to native space using FMRIB’s Linear Image Registration Tool (FLIRT) and nearest neighbor interpolation, then thresholded to zero-out the bottom 10% of *z*-statistics. After alignment to native space, we ran clustering over the map and excluded voxels that did not belong to a cluster of at least 10 voxels, resulting in a mean number of 1,354 voxels per participant (range = 1, 058 *−* 1, 599 voxels).

### fMRI experiment design

The full study comprised a total of 4 or 5 sessions per participant. Four functional runs were collected during each session and participants were offered a break between runs. Sessions were scheduled on separate days with a target of 24 hours between sessions and a maximum of 48 hours between neurofeedback sessions (mean = 27.24 hours, range = 13 *−* 48). All sessions had to be completed within 7 days.

#### Joystick session

In Session 1, participants practiced the video game task by navigating the avatar with a MR-compatible joystick (Current Designs Tethyx). The joystick had a two-axis (X, Y) shaft controller, allowing an angular range of 30^*°*^ (*±*15^*°*^) and a square range in X and Y movement planes of *±*15^*°*^. Accordingly, participants could move the avatar forward at a constant speed and rotate it at any angle (*−*180^*°*^ to 180^*°*^). Upon release, an internal spring forces the joystick shaft back to its vertical position. The joystick was connected to the experimental computer via a fiber optic cable and interfaced directly with the Unity engine like a standard gaming joystick.

Participants held the joystick with their dominant hand and placed the base either on the scanner bed at their dominant side or on their abdomen, securing the base with their non-dominant hand. Once situated in the scanner, participants were allowed up to two practice trials before the first scan to make sure they were comfortable holding and manipulating the joystick without moving. Participants were instructed to navigate the avatar from its starting location to the target within the time allowed (60 s in the first run, +10 s for each subsequent run) while avoiding obstacles. They were informed that they could explore the environment (i.e., not head directly along the display line to the goal), which allowed us to broadly sample the state space of their brain activity across movement patterns and arena locations. The resulting duration and number of trials for each run, session, and participant thus varied (mean = 19.1 trials, range = 8 *−* 26 trials). A trial ended upon reaching the flag and the game paused for 6 s (3 TRs) before the next trial began.

#### Neurofeedback calibration

Data collected during the joystick session were used to initialize the manifold-based neurofeedback procedure used in subsequent sessions. fMRI data processing used fMRI Expert Analysis Tool (FEAT) version 6.00 [63], part of fMRIB’s Software Library (FSL) version 6.0.5. EPI and anatomical images were skull-stripped using the Brain Extraction Tool (BET) [64]. Susceptibility-induced distortions were measured via the opposing-phase spin echo volumes and corrected using FSL’s topup [65]. Each functional run was high-pass filtered with a 100 s period cutoff, corrected for head motion with Motion Correction using FLIRT (MCFLIRT) [66], corrected for slice timing, and smoothed spatially with a Gaussian kernel (5-mm full-width half-maximum; FWHM). Then, functional images were registered to the participant’s T1-weighted anatomical scan using boundary-based registration (BBR) [67] and to a 2-mm MNI standard brain using 12 degrees of freedom.

Information about the state of the game on each trial, including the avatar’s location in the arena over time, was collected at 60 Hz and saved from Unity. This information was processed with custom Python scripts, downsampled to 0.5 Hz using average pooling to match the temporal resolution of the fMRI acquisitions, and shifted 2 samples (4 s) to correct for the hemodynamic lag in the corresponding fMRI data.

fMRI data samples from the joystick-based navigation trials (the timepoints between trials were excluded) were then divided into training (80%) and testing (20%) sets. The training set was used to fit a 20-dimensional T-PHATE embedding [40]. We defined three latent components of the T-PHATE manifold: the eigenvectors associated with the greatest, second greatest, and smallest eigenvalues. These components *C* correspond with the “intuitive mapping” (IM; *C*_*IM*_), the “within-manifold perturbation” (WMP; *C*_*W MP*_), and the “outside-manifold perturbation” (OMP; *C*_*OMP*_) components used for neurofeedback. Finally, we trained a MRAE with two training objectives: to reconstruct the input fMRI data in voxel space, and to learn a latent representation that closely resembles the data’s ground-truth T-PHATE manifold geometry. This training allows the MRAE to extend the T-PHATE manifold to incoming data points collected in real-time. All T-PHATE, MRAE training, and component determination was performed on the training set of the joystick session data (80%). Detailed technical explanations of these steps are provided in the “Neural manifold learning” section below.

After the autoencoder was trained on the training set, its weights were frozen and the testing set was passed through the encoder *f*, extracting its embedding in the T-PHATE manifold. We then projected the embedded data onto *C*_*IM*_, *C*_*W P*_, and *C*_*OMP*_ as previously defined, to measure the distribution of untrained data along the components of the manifold. The 1st and 99th percentiles of the distributions of untrained data along each component were fixed to represent turning *−*180^*°*^ and 180^*°*^ in the game, respectively.

#### Neurofeedback sessions

After the joystick session, participants returned to the scanner for three or four fMRI neurofeedback sessions. During each neurofeedback session, participants completed four functional runs (*∼* 10 minutes each) during which their task was to use their brain to make the avatar walk to the flag as directly as possible. Specifically, the instructions were to “generate a mental state that made the avatar walk to the flag.” We emphasized to participants that there was no single right strategy to use to accomplish this task; they would need to try a variety of strategies and possibly switch between strategies. Indeed, participants reported a wide range of strategies in a post-study questionnaire (Supplemental Table S1). Participants were reminded at the start of each run that the avatar’s movement was a reflection of their brain states over the past 4–6 s, and that the avatar’s movement would be a delayed reaction to their thoughts.

We outlined the staircasing training procedure to participants as followed. Participants were told that there were “bumpers” (akin to those on a bowling lane) providing assistance in keeping the avatar walking relatively straight to the goal at first (i.e., at “lower levels”). As brain activity resulted in more accurate movement, participants would “level up” and the bumpers would go down. Thus, the avatar could veer farther off-course and require brain activity to be more accurate in order to go straight. Participants were instructed that their goal was to continue “leveling up.” The first neurofeedback run started at level 0, and at the end of each trial, participants were presented with a feedback display explaining performance as the percentage of distance traveled over the shortest possible path to the goal. They were then informed if this error was higher, lower, or equal to the average error of their previous three trials. The staircasing procedure began after the third trial. Participants “leveled up” if their error decreased, “leveled down” if error increased, or stayed the same if their error matched prior trials.

At the start of each neurofeedback run, there was a 20 s (10 TR) delay before the first trial. Task instructions were presented during this time, reminding participants that their goal was to get to the flag as quickly as possible and to remember that there would be a delay between their brain state and the avatar’s movement. Then the first trial began and participants completed as many trials as they could in approximately 10 minutes (300 TRs), with a 6-s break between trials. After that, the scan was ended manually by the experimenter and participants rested until the next run. The first trial of the next run began at the final “level” of the previous run.

#### Real-time data processing

Every 2 s, a whole-brain volume was acquired as a DICOM image and sent via fiber network from the local Siemens scanner console to a remote, HIPAA-aligned high-performance compute cluster. The first volume of each run *V*_1_ was smoothed with a 5-mm FWHM Gaussian kernel to match the preprocessed data from the joystick session and saved as that run’s reference volume, to which all subsequent volumes were aligned. FLIRT [66] was used to align *V*_1_ to *V*_*ref*_, a reference mean functional image from the joystick session. The resulting transformation matrix *T* served to transform each subsequent volume to *V*_*ref*_, the space in which the ROI mask of navigation-related brain voxels was defined. Each volume *V*_*t*_ was smoothed, motion corrected with MCFLIRT to *V*_1_, aligned to *V*_*ref*_ via *T*, and masked to extract voxels in the feedback ROI.

The first 10 brain volumes in each run were used to estimate mean and standard deviation parameters for each voxel. These parameters were then used to normalize (i.e., *z*-score) the activity of each voxel in the 11th volume and updated with each time step to reflect prior volumes. That is, for a given voxel’s activation vector *x* consisting of *t* timepoints, its normalized activity at time *t, v*_*t*_, would be determined by Equation 1:

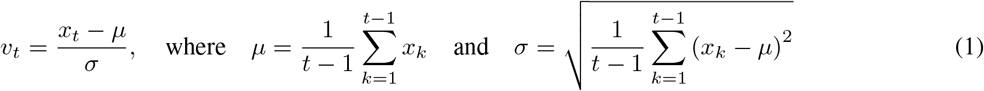

After selecting voxels from the navigation mask and normalizing them, the data *V*_*t*_ were passed through the trained MRAE encoder *f*, such that *f* (*V*_*t*_) yielded the embedding of *V*_*t*_ onto the T-PHATE manifold. Finally, *f* (*V*_*t*_) *· C*_*ses*_ mapped the T-PHATE-embedded data onto the feedback component for that session *C*_*ses*_, which determined the angle of movement *α ∈* (*−*180^*°*^, 180^*°*^). The value of *α* was transmitted back to the scanner presentation computer via the fiber network link to a script running PsychoPy, which was time-locked in communication with the Unity video game via a TCP connection and used to determine the next step, *γ*, such that:

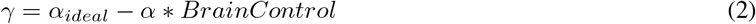

where *α*_*ideal*_ was the angle that would keep the avatar along the straightest path to the goal and *Brain Control* was the proportion of control *α* had over the avatar’s movement.

At the end of each trial *T*, the error *ϵ*_*T*_ was computed as the ratio of the total distance traveled by the avatar relative to the shortest possible path between the start and goal locations. *Brain Control* was then staircased depending upon the *ϵ*_*T*_ and *ϵ*_*prior*_ (defined as the average error over the prior two trials) [47], such that the *Brain Control* at trial *T* + 1 was determined by Equation 3:

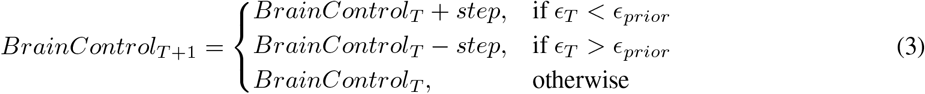

Participants were given feedback along the intuitive component for the four runs of the second session (first neuro-feedback session). *Brain Control* did not plateau during this session for one participant and so they repeated the IM session in their second visit and completed five sessions total instead of four. For subsequent neurofeedback sessions, participants began the first trial of the first run with *Brain Control* set to the final *Brain Control* value from the end of their IM session plus 20%. They began with one run of IM training to re-calibrate the feedback procedure before receiving the within-manifold or outside-manifold perturbations in the second run.

#### Simulated null participants

To validate that the BCI learning effects were not a byproduct of our feedback pipeline or staircasing procedure, we simulated 20 null participants using the *fmrisim* module of the BrainIAK package [46]. Each was based on the brain mask and noise parameters of a real participant, including their signal-to-noise ratio (SNR), signal-to-fluctuating-noise ration (SFNR), and FWHM. Using these noise parameters derived from the real fMRI data, fmrisim generated system and temporal noise sources (i.e., physiological, drift, autoregressive), resulting in noise timecourses for 300 timepoints for each voxel in the brain mask. This was repeated for 16 times for each of the 20 simulated participants, to match the 4 runs x 4 sessions per real participant.

With each simulated null participant, we repeated the T-PHATE and MRAE training procedures from the first joystick session (Figure 2C and Figure S2). For each subsequent session, we ran the neurofeedback protocol over the simulated brains by writing out each image as a DICOM every 2 s, processing it and mapping it to a direction of movement, and returning that to the Unity display, where the accuracy was calculated and *Brain Control* adjusted as with the real participants. We then tested within- and outside-manifold perturbations in a counterbalanced order, as for real participants.

### Neural manifold learning

A key development introduced in this study is the use of diffusion geometry methods [37–39] to perform neural manifold learning and define an intrinsic manifold of neural activity to target with neurofeedback. Prior studies using low-dimensional data representations as input for BCIs have relied on linear dimensionality reduction methods (e.g., PCA, factor analysis) to approximate the intrinsic manifold of brain activity as a linear subspace. More recent studies have shown that high-dimensional neural population activity can be summarized within a lower-dimensional nonlinear manifold [40, 68, 69]. Thus, the linear approximation of a nonlinear manifold in prior work may not fully capture the intrinsic manifold of brain activity and what activity patterns are on versus outside the manifold (Figure S6).

fMRI signals have high noise in both the space and time dimensions, so extracting behaviorally meaningful brain activity from fMRI often requires aggregating activity across many trials or participants, or reducing dimensionality via averaging, to improve signal-to-noise. Recently, we have applied manifold learning to represent fMRI activity from single participants during cognitively complex tasks [40]. We designed a novel three-step procedure to learn and apply nonlinear manifolds in real-time fMRI: (1) learning the intrinsic manifold of single-participant fMRI activity via T-PHATE, which builds upon the classic diffusion maps algorithm [37]; (2) training a manifold-regularized autoencoder (MRAE) to quickly embed new brain volumes onto the learned T-PHATE manifold and projecting them to the BCI mappings; and (3) identifying within- and outside-manifold components based on the principal components of the MRAE latent space as BCI mappings. These three steps are elaborated upon below.

### T-PHATE

The T-PHATE algorithm is a dual diffusion-based manifold learning method for discovering data geometry and latent dynamics from complex, biological timeseries data. We used T-PHATE to learn a low-dimensional manifold of brain activity in our study because T-PHATE has been demonstrated to faithfully capture cognitively meaningful signals and task information in single-participant fMRI data [40]. T-PHATE takes as input multi-voxel activity patterns (i.e., a matrix with timepoints/samples as rows and voxels/features as columns) and learns two “views” among pairs of samples: a PHATE-based [39] affinity matrix and a temporal autocorrelation-based affinity matrix. Extending upon the classic diffusion maps algorithm [37], PHATE provides an accurate, de-noised representation of local similarities (via an *α*-decay kernel) and global relationships (via a diffusion potential distance), without many assumptions about a hypothesized manifold structure [39]. The autocorrelation matrix models the temporal dynamics across data samples by computing the correlation of each voxel’s timeseries with lagged versions of itself. This kernel captures both the temporal dynamics related to the measured BOLD signal and those related to the temporally diffuse cognitive processes occurring within a given multi-voxel pattern. The PHATE and autocorrelation views are converted into transition probability matrices and then combined with alternating diffusion, before embedding into an *M* -dimensional representation using metric multi-dimensional scaling (m-MDS). T-PHATE embeddings were performed for individual participants. fMRI timeseries data input to the T-PHATE algorithm were masked to include only voxels in the navigation mask, *z*-scored within voxel and fMRI run, and concatenated across runs.

### MRAE

T-PHATE embeddings improved access to task-relevant information from the brain data, as shown by the improvements in location decoding and arena representation (Figure 1C,D). A key limitation of manifold learning methods such as T-PHATE is that they are not readily extensible to new data samples. To avoid needing to re-fit the T-PHATE algorithm entirely with each incoming fMRI volume in real-time (which would be prohibitively slow), we trained a MRAE to (1) reconstruct the voxel-resolution fMRI data and (2) learn the correspondence between input fMRI data and the corresponding points on the T-PHATE manifold [44, 45]. This operation was trained via manifold regularization penalty, which minimizes a geometric loss function of the distance between fMRI samples in the autoencoder’s bottleneck and the initial T-PHATE embedding. The encoder thus learned a nonlinear mapping between the fMRI data in voxel space and the fMRI data in T-PHATE space; after training was complete, we could use the encoder to embed new fMRI samples onto the T-PHATE manifold and it faithfully interpolated along that manifold.

The MRAE had three fully connected layers in each the encoder (*f*) and decoder (*g*), with a bottleneck latent space layer in between. As input, we gave the model both the training data *Y* (in this experiment, 80% of the data from the joystick session) and the T-PHATE embedding of the training data (*E*(*Y*)). The input layer to *f* and output layer of *g* had *n*_*voxels* units, which was participant-dependent (i.e., the number of voxels in an individual’s navigation network mask). Hidden layers had 256 – 128 – 64 – 20 – 64 – 128 – 256 units, respectively, and leaky ReLU activations were applied on all layers. We used the Adam optimizer, batch size of 64, and learning rate of 0.001. The model was trained with a reconstruction error penalty for each sample seen at training time, to minimize the mean-squared error between *Y* and Ŷ, or *f* (*g*(*Y*)). Thus, given participant’s data *Y*, where *Y* has *k* timepoints, encoder *f*, and decoder *g*, the reconstruction penalty was defined by Equation 4:

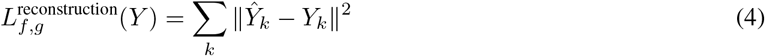

The model was also trained with a manifold regularization, which pushed the bottleneck layer to have the same manifold geometry as the initial T-PHATE embedding of *Y, E*. This was computed by Equation 5:

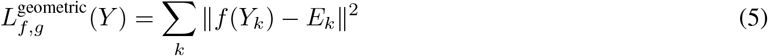

To combine *L*^geometric^ and *L*^reconstruction^, we used a coefficient *γ* which controlled the amount of geometric regularization to the hidden layer of the MRAE. The combined loss *L* thus became Equation 6:

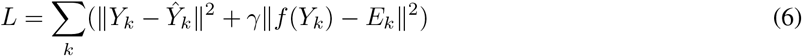

which was optimized over encoder *f* and decoder *g*. After training, these weights were frozen and the testing set of data from the joystick task (the remaining 20% of data) were passed through *f* to obtain their embeddings in the latent space.

### Defining manifold components

We used manifold components to map the data from the T-PHATE manifold to the avatar’s movement. We learned these components using PCA over the MRAE latent space (post-training), which yielded the eigenvectors of the T-PHATE covariance matrix. We selected the three eigenvectors which captured the greatest, second greatest, and least variance in the manifold-embedded brain activity. We validated that this approach of using the T-PHATE-based eigenvectors faithfully reflected the nonlinear latent structure of noisy data (Supplemental Figure S6). Experimentally, we refer to these eigenvectors as the neurofeedback mapping components, and we take the first and second components to be *C*_*IM*_ and *C*_*W MP*_ and the 20th component to be *C*_*OMP*_. These components were fitted individually on each participant’s training data from the joystick session and held constant through the experiment (i.e., not re-fitted on subsequent days).

## Data analysis

### Joystick session validation

We validated that the T-PHATE embeddings captured task-relevant signals during the joystick session. Qualitatively, we visualized the first three dimensions of the T-PHATE embedding timeseries and colored each point as the avatar’s coordinate in the game arena at that timepoint. We depict five participants’ embeddings of four consecutive trials, which visually show clustering according to game arena coordinate, such that similarly-colored points are clustered together despite being collected during different trials (Figure 1B).

We next attempted to decode the avatar’s coordinates from the manifold embeddings. We trained two multiple linear regression models, one to predict X coordinates and one to predict Y coordinates, from the T-PHATE embedding timeseries in the navigation network, using leave-one-run-out cross-validation. On the held-out run, models were scored as the Euclidean distance between the model-predicted X/Y coordinates and the true X/Y coordinates. To assess the benefit of T-PHATE, we performed the same analysis using the voxel-resolution data (*∼* 1, 300 voxels) prior to running T-PHATE (Figure 1C).

We also evaluated the representational similarity of brain activity and arena locations. In other words, are more proximal locations in the arena represented more similarly in the brain than more distant locations? We quantified this similarity by extracting fMRI activity patterns for each location and calculating the Pearson correlation of all pairs of locations to populate a location-by-location correlation matrix. We generated a distance matrix for the arena by populating another location-by-location matrix with the pairwise distance in space between each pair of locations. We then tested for representational similarity between the brain’s correlation matrix, calculated from both voxel-resolution and T-PHATE embeddings, and the arena’s distance matrix using a Mantel test [42, 70], which computed the statistical significance of the Spearman correlation between matrices with a randomization procedure (10,000 iterations). The Mantel test was performed for each participant and converted to a *z*-score (Figure 1D).

### BCI learning during neurofeedback

The staircased *Brain Control* parameter scaled with task accuracy (i.e., stepping up with better performance, down with worse performance), so Δ*BrainControl* serves as a metric of BCI learning in each session. Across trials, we tested whether Δ*BC* was significantly greater than zero using permutation testing with cluster-based correction. As a baseline, we computed the changes in *Brain Control* for each session type using the simulated null participants.

### Neural realignment with BCI learning

To quantify the neural changes underlying BCI learning, we computed the proportion of neural variance explained by each component of the manifold. Namely, using the 20-D T-PHATE embedding of each fMRI timepoint (excluding rest between trials) (*f* (*V*_*t*_)), we computed the variance explained by each eigenvector. We divided this value by the overall variance of the data and multiplied by 100 to get the percentage of explained variance (*PEV*) along each component. We focused on the *PEV* along the IM (*f* (*V*_*t*_) *· C*_*IM*_), WMP (*f* (*V*_*t*_) *· C*_*W MP*_), and OMP (*f* (*V*_*t*_) *· C*_*OMP*_) components. We then took the final run’s *PEV* and subtracted it from the first run’s *PEV* to get Δ*PEV*, our measure of neural realignment. In the main analyses (Figure 4), we report Δ*PEV* for each component in its corresponding session (i.e., Δ*PEV* for the IM component during the IM session). Here we report Δ*PEV* for each component in non-corresponding sessions (i.e., Δ*PEV* for the IM component during the WMP session) to show the specificity of the observed neural learning effects (Supplemental Figure S4).

### Whole-brain decoding

The navigation mask of brain regions used for neurofeedback showed neural realignment, but changes could also be reflected elsewhere in the brain. Using a whole-brain searchlight analysis, we computed the accuracy of decoding the avatar’s coordinates in the game arena from multivoxel patterns of fMRI data during the first and final runs of neurofeedback for each session type. Searchlights were centered on every voxel in the brain, each surrounded by a sphere with a radius of 3 voxels. For each searchlight, we trained a linear model (ridge regression, himalaya package in Python; [71]) to predict the avatar’s location (X, Y coordinates) at each timepoint from the fMRI activity pattern across voxels in that searchlight using leave-one-trial-out cross-validation. Models were scored based on the Euclidean distance between the predicted and true coordinates in the held-out trial, then averaged across folds to get one spatial map of distance scores for the first and final neurofeedback runs of each session type (lower distance means improved decoding). We subtracted the final run’s distance map from the first run’s distance map, then normalized the difference by the sum of the first and final run’s distances. This results in a value between -1 and 1 for each voxel, with 1 indicating a larger relative distance in the first run than the final run. Finally, we tested the statistical significance of these difference scores in each session type using FSL’s randomise function [72] with variance smoothing and a 5-mm sigma, corrected for multiple comparisons with threshold-free cluster enhancement (TFCE) [73]. Note that these analyses were, by necessity, performed in voxel space, as the searchlights were different sets of voxels than those used to define the T-PHATE manifold.

### Statistical analyses

We used nonparametric tests throughout to determine statistical significance. Participant-level bootstrap resampling (10,000 resamples) was used to assess random-effects reliability across participants for comparisons of variables against chance or between conditions [74]. One-sided tests were used for directional hypotheses and two-sided tests for nondirectional hypotheses. The significance of effects at the trial level were evaluated using permutation tests, with cluster-based multiple comparisons correction across adjacent trials. To evaluate individual differences in brain-behavioral relationships, we used a linear mixed model (estimated with REML and nloptwrap optimizer) including participant and session number as random effects. 95% Confidence intervals and *P* -values were computed using a Wald t-distribution approximation.

## Data availability

Raw and preprocessed functional and anatomical images and behavioral data will be released publicly upon acceptance.

## Code availability

The task code was programmed with C# for Unity (unity3d.com; Version 2019.4.12f1) and is available at: github.com/ericabusch/avatarRT_task. The real-time fMRI experiment used the open-source rtCloud frame-work (rt-cloud.readthedocs.io) [43]. Our experiment, preprocessing, and analysis scripts are available at: github.com/ericabusch/avatarRT_analysis.

## Author contributions

E.L.B., S.K., and N.B.T-B. conceived the study. E.L.B, G.L., S.K., and N.B.T-B. designed the experiment. E.L.B. and E.C.F. conducted the neurofeedback experiment and the simulation experiments. E.L.B. analyzed the data with support from E.C.F. and feedback from all authors. E.L.B., S.K., and N.B.T-B. contributed to the development of data processing algorithms. E.L.B. and N.B.T-B. wrote the manuscript with extensive feedback from all authors.

## Acknowledgements

E.L.B. was supported by a NSF Graduate Research Fellowship (award no. 2139841). G.L. was supported by Canada CIFAR AI Chair and Canada Research Chair in Neural Computations and Interfacing. S.K. was supported by the NIH (grant nos. R01GM135929 and R01GM130847), an NSF Career Grant (grant no. 2047856) and a Sloan Fellowship (grant no. FG-2021-15883). N.B.T-B. was supported by an NSF Grant CCF (grant no. 1839308) and CIFAR. The authors would like to thank Lillian Behm, Thomas Botch, May Conley, Gloria Feng, Joseph Heffner, Chang-Hao Kao, Ariadne Letrou, Kailong Peng, Juliana Trach, Tristan Yates, Xihan Zhang, Irene Zhou, and Yannan Zhu for help with data collection.

## Supplementary information

**Figure S1:**
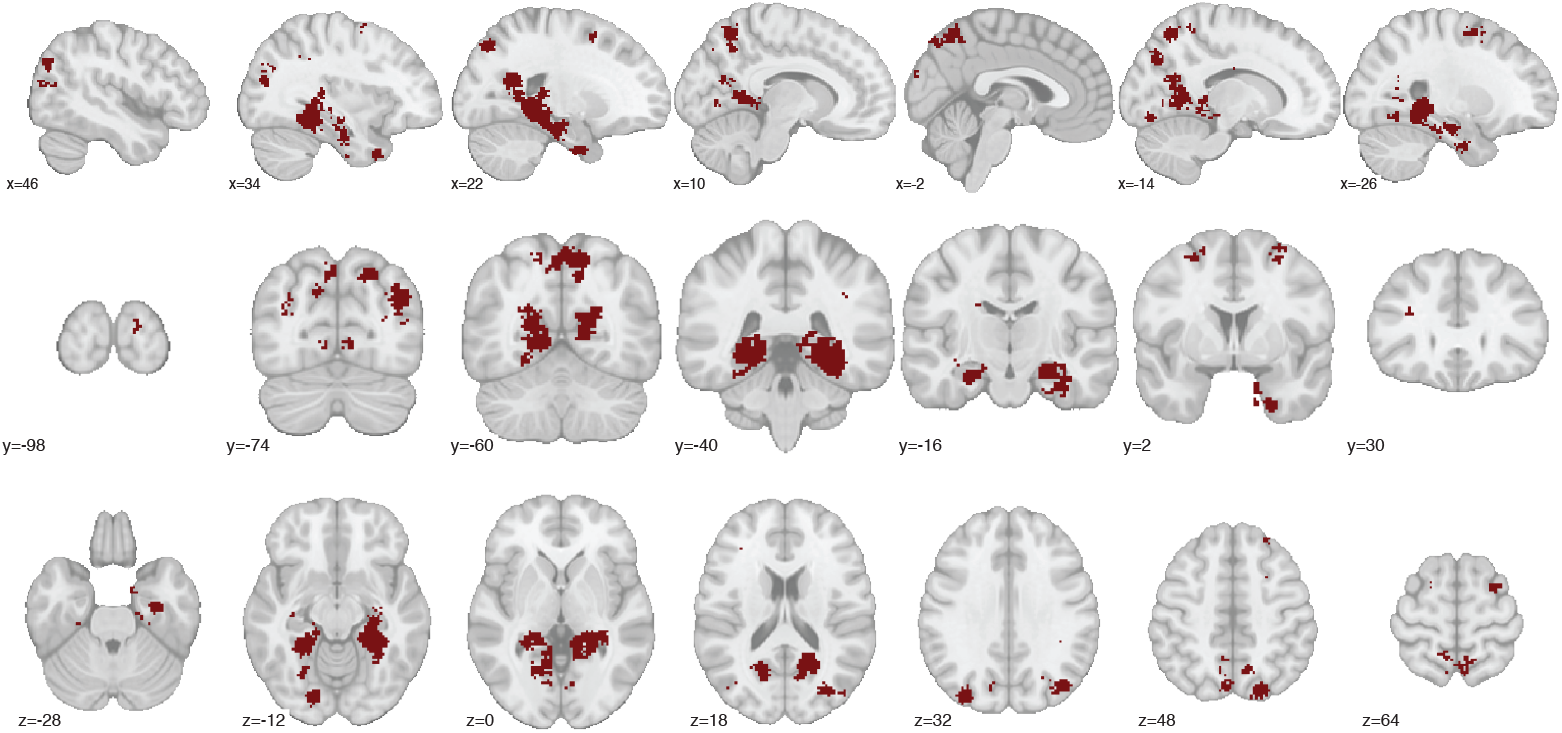
Navigation network targeted with neurofeedback. Sagittal, coronal, and axial slices of the MNI template brain showing voxels which were used for neurofeedback. Primary regions include hippocampus, medial temporal lobe cortex, precuneus, lateral occipital cortex, and parahippocampal cortex.

**Figure S2:**
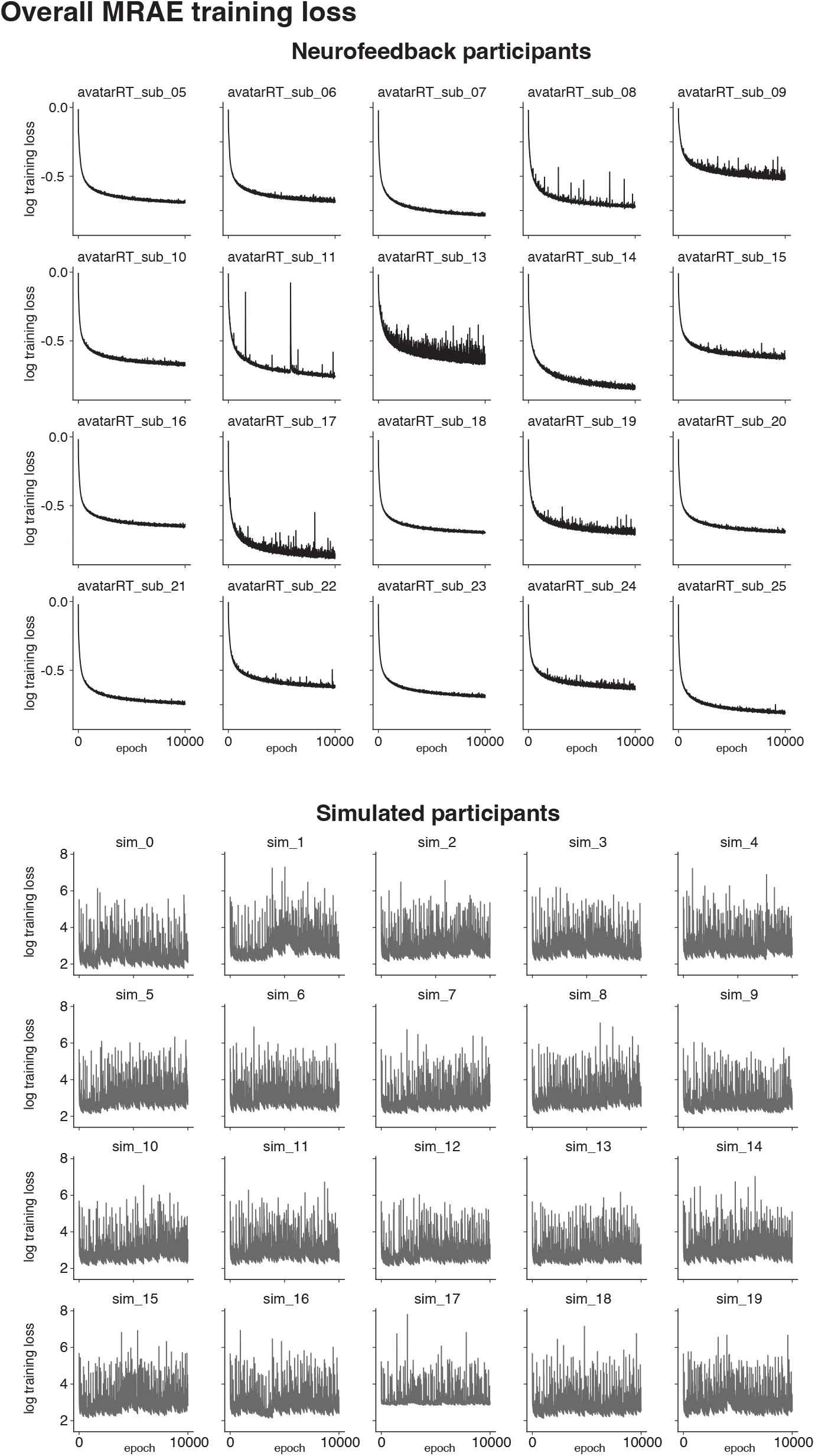
Participant-level MRAE training loss. MRAE training loss curves (*L*) of 10,000 training epochs for real participants (top, black) and simulated null participants (bottom, gray). See Figure 2C for averages. Overall loss is the weighted combination of the reconstruction error and the manifold regularization penalty. MRAE training loss decreased rapidly and plateaued for all real participants by *∼* 5, 000 epochs, though final training loss plateaued at different points across participants. As expected, simulated null participants showed no decrease or convergence (note the substantially higher y-axis range).

**Figure S3:**
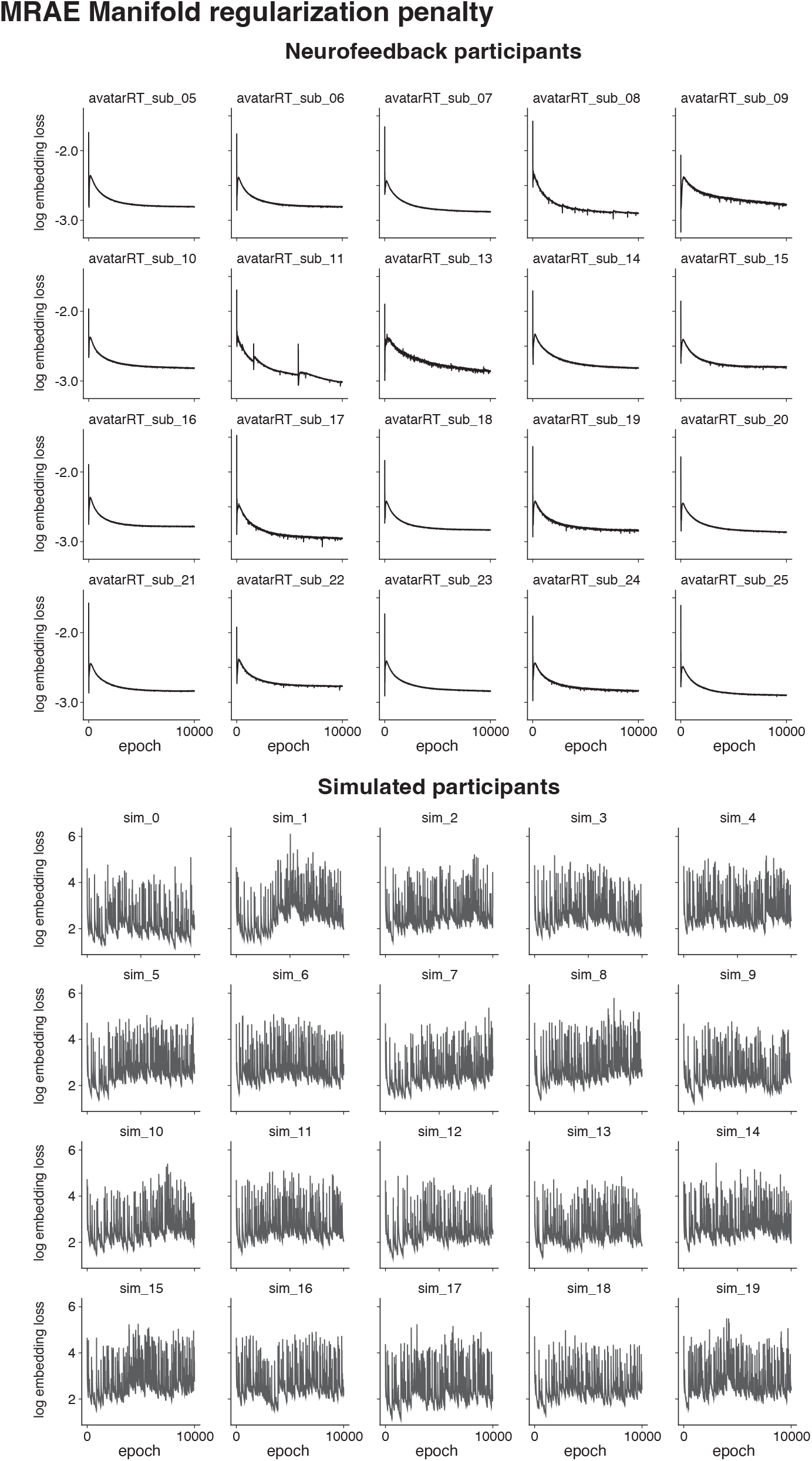
Participant-level manifold regularization penalty. Manifold regularization penalty curves (*L*^geometric^) of 10,000 training epochs for real participants (top, black) and simulated null participants (bottom, gray). See Figure 2D for averages. The decreasing values for real participants indicate that the encoder *f* learned weights which successfully embedded new samples into their respective positions along the T-PHATE manifold, meaning that there was meaningful, task-related, and predictable structure in the input data that matched the T-PHATE representation. This was not possible for the simulated null participants, who showed no decrease or convergence (note the substantially higher y-axis).

**Figure S4:**
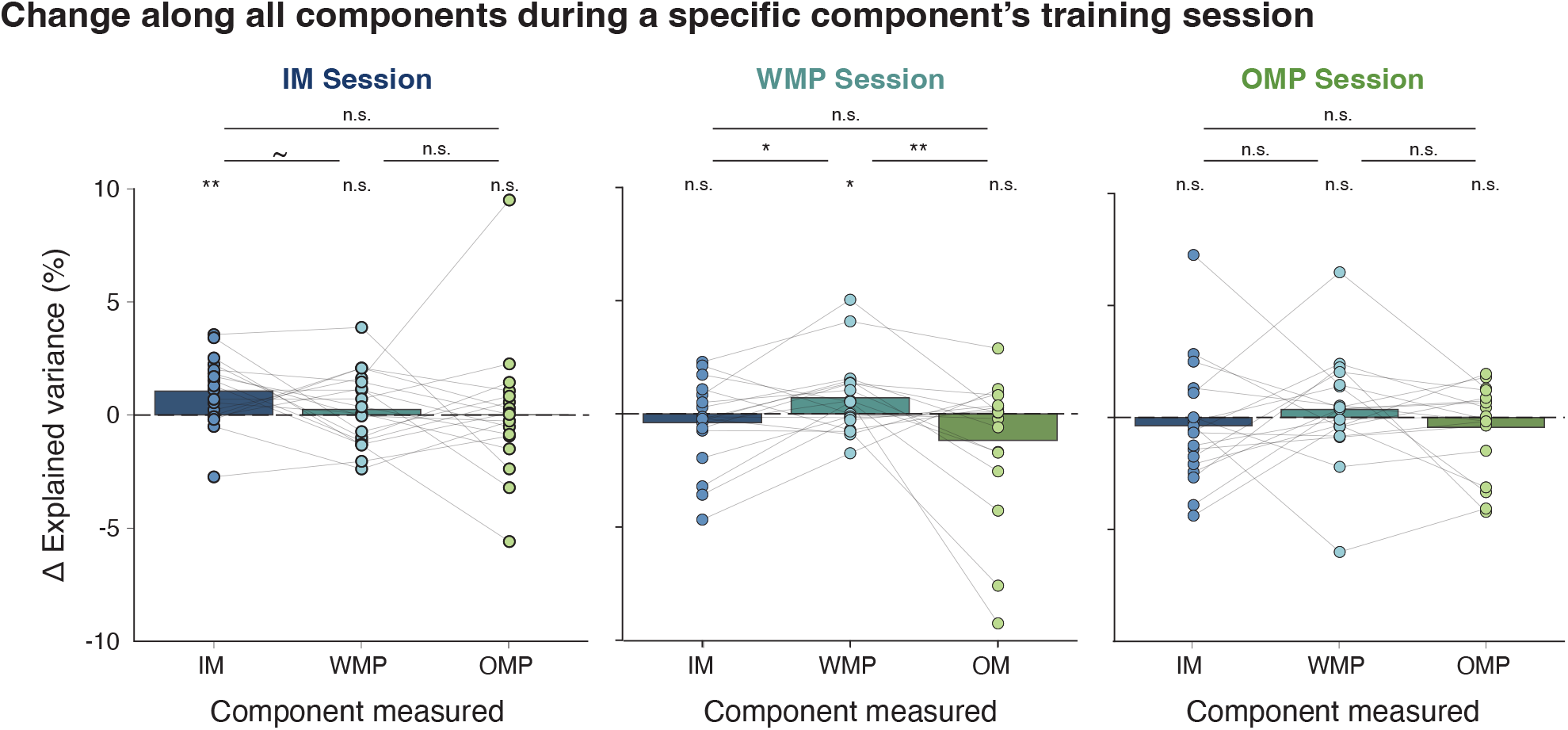
Neural realignment with all feedback components. We measured the change in the percentage of explained variance (Δ*PEV*) along the IM, WMP, and OMP components over the course of training for each component. That is, for the IM training session, we measured the Δ*PEV* for the IM, WMP, and OMP components, though the WMP and OMP components were not being trained during that session. Neural realignment was specific to the component being trained, as IM and WMP showed increased Δ*PEV* only when they were being trained in the IM and WMP sessions, respectively; as expected, none of the components showed increased Δ*PEV* in the OMP session in which learning did not occur.

**Figure S5:**
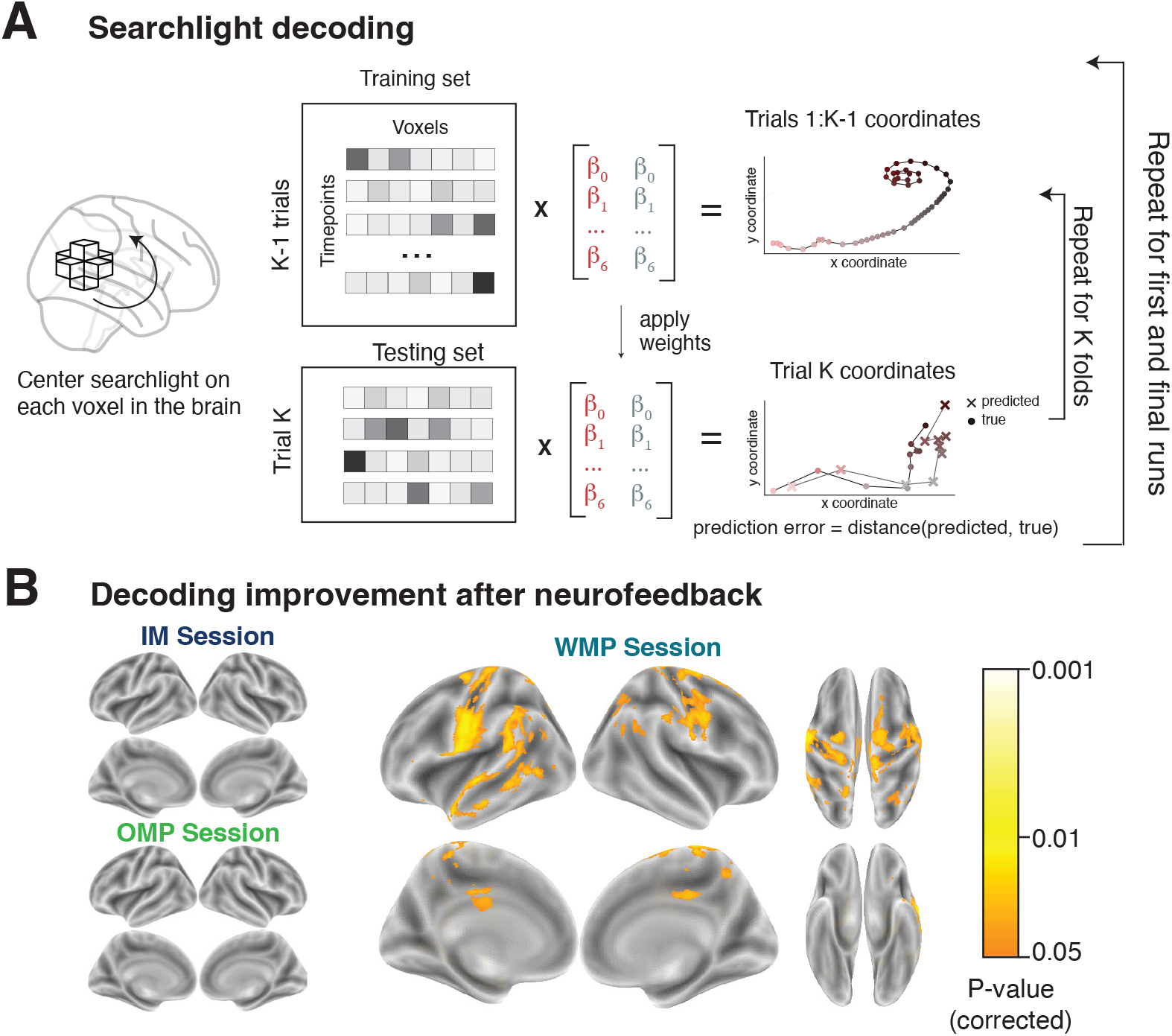
Whole-brain improvements in task decoding. (A) BCI learning can alter non-target brain regions, such as those involved in cognitive strategies and modulation of target regions [59, 75–77]. An exploratory searchlight analysis was used to test for improved task representation in regions across the whole brain, by decoding the avatar’s coordinates in the game arena from each timepoint using ridge regression with nested cross-validation. Regression models were trained and tested with leave-one-trial-out cross-validation in the outer loop, and hyperparameter optimization was performed in the inner loop. Regression models were scored as the Euclidean distance between model-predicted arena locations and true participant locations. We measured decoding improvement from the map of prediction errors across the brain in the first minus last runs of a session; lower error means improved prediction and thus a clearer representations of the task. (B) Voxelwise map for each session type depicting searchlights with improved location decoding (TFCE corrected). No regions showed improved decoding accuracy in the IM and OMP sessions. This was expected for OMP given that there were no indicators of BCI learning for this session type. In the IM session, neural realignment was not found to predict BCI learning, suggesting that this component was already controllable; this was reflected by the lack of decoding improvement, which indicates that strategies and modulatory patterns do not need to change from their original activity to support task performance. In the WMP session, decoding accuracy of arena location improved across neurofeedback runs in primary motor cortex (notably areas related to hand and finger movement) as well as in parts of the executive control, dorsal and ventral attention networks. Only 195 of the 26,179 voxels with significantly greater (*P <* 0.05, corrected) decoding accuracy after BCI learning in the WMP session overlapped with the voxels included in the navigation mask (i.e., 3.2% of voxels in the mask).

**Figure S6:**
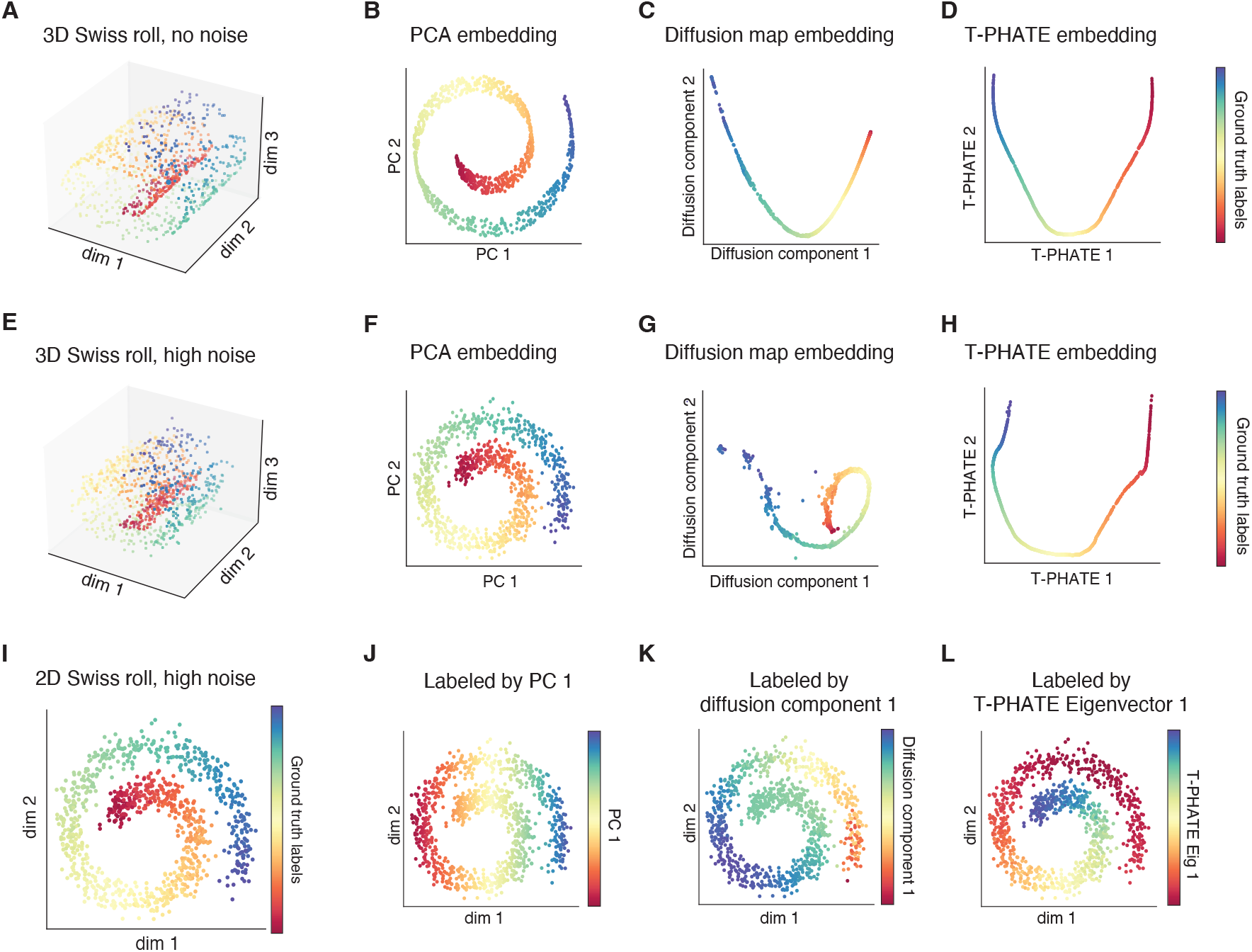
Nonlinear methods improve detection of on-manifold activity in nonlinear data. (A) Simulated, noise-free Swiss roll dataset of 1,000 points displayed in ambient 3-D space (standard deviation of Gaussian noise, SD = 0.0). Data colors represent position along the main dimension of the ground-truth manifold. (B–D) 2-D embeddings of (A) with (B) PCA, (C) diffusion maps [37], and (D) T-PHATE [40]. Nonlinear dimensions (i.e., diffusion maps and T-PHATE) reveal an “unrolled” representation of the Swiss roll data, representative of its structure. (E) Swiss roll dataset with high noise (SD = 1.0). (F–H) Embeddings of (E) as in (B–D). With higher noise added to the 3-D data, T-PHATE remains able to unroll the true data manifold, whereas the diffusion map remains slightly coiled. (I) Swiss roll with high noise in a 2-D ambient space, colored by its ground-truth manifold positions. (J–L) Data as in (I), now colored by each point’s loading onto the first component of the embeddings from (J) PCA, (K) diffusion maps, and (L) T-PHATE. As in the ground truth labeling (I), the loadings shown in (L) track the coil of the Swiss roll, whereas (J, K) do not.

**Table S1:** Strategies reported by participants (*N* = 18) in debriefing questionnaire administered after completion of all sessions. This question was phrased as: “What strategies di7d you try in order to control the avatar? How successful did you think these strategies were?” and participants typed their answers into a blank text box. Responses here are lightly edited for clarity and conciseness.

| Participant | Self-reported strategies |
| --- | --- |
| 5 | (1) Thinking about moving straight, (2) thinking about following the yellow line, (3) following the diagonal path along the green check marks, (4) thinking about heading directly to the flagpole (most successful). |
| 6 | (1) Thinking about nice things (vacations, dates, etc), (2) doing arithmetic, (3) planning for the next months, (4) looking at the flag, (5) looking at the avatar's feet (most successful). |
| 7 | Various strategies, none of which were obviously successful. |
| 8 | (1) Thinking about happy, sad, stressful, and pleasurable memories ((pleasurable memories were more successful than non-pleasurable), (2) replaying movies that they had seen multiple times and could recall vivid visual and auditory features (repeating the same movie was less effective the second time), (3) thinking about the octopus on the flag (successful some of the time). |
| 10 | (1) Thinking about walking, (2) thinking about playing games, (3) thinking about playing sports, (4) imagining moving toward the flag, (4) focusing on the scenery and trees, (5) reminiscing old memories, (6) trying to embody happiness, sadness, or other emotions. |
| 11 | Playing a song in my head in Swahili. If the song was in Swahili, the avatar went straight to the flag, but if I accidentally mentally sang songs in English, the avatar spun. I also instructed the avatar at the same time, but the instructions had to be in English while the song was still in Swahili. This was very successful but if I scrambled my languages, it took a while to get working again; overall, it was easy to overthink and get confused. |
| 13 | (1) Thinking "straight" to keep the avatar on the straight path (most successful), (2) keeping my eyes on the goal location, (3) envisioning a sprinter running down the final stretch of a race, (4) envisioning myself walking, (5) reminding myself to not get distracted, (6) thinking "right" or "left" to get back on the path when the avatar veered off course. |
| 14 | (1) Singing to myself (not successful), (2) thinking happy thoughts (somewhat successful), (3) pretending I was holding the controller (somewhat successful), (4) pretending I was walking to a destination while imagining and creating a mental scene (most successful). |
| 15 | (1) Looking for patterns in the squares ahead, (2) imagining I was walking my dog and he was pulling me toward the flag because there was food or other dogs over there (most successful but required very vivid imagery). |
| 16 | (1) Following target with eyes and repeating "octopus flag" over and over (successful in the beginning then stopped), (2) following the target with eyes and thinking about daily life (most successful), (3) imagining moving the joystick with hand (sometimes successful), (4) grinding teeth when going off path (painful and not successful). |
| 17 | (1) Telling the avatar mentally to move, (2) relaxing my mind and eyes by focusing on the sound of the MRI and trying not to get stressed out by going in the wrong direction (most successful). |
| 18 | (1) Thinking "forward" over and over (somewhat successful initially at low levels), (2) thinking about my mind moving the avatar to the target, (3) staring intensely at the flag (most successful). |
| 19 | (1) Thinking "left" and "right" (most successful), (2) thinking about tapping on my leg. |
| 21 | (1) Imagining running to break up a fight between a pair of kids before one gets bitten, (2) thinking about walking to work, (3) mentally running through my to-do list, (4) relaxing and counting the avatar's steps, either to get to the flag or 1 to 10 on repeat (most successful). |
| 22 | Thinking about cooking different meals from start to finish. |
| 23 | (1) Imagining scenarios that involved forward, one-directional movement like dog walking, (2) imagining I was surfing or swimming (most successful). |
| 24 | (1) Pretending that someone or something I wanted to reach was at the flag, (2) imagining I was walking down the street to my house (both were successful). |
| 25 | (1) Mentally singing "Just Keep Swimming" from *Finding Nemo*, (2) imagining walking to class, (3) intentionally not focusing on the screen and trying not to overthink my performance (most successful). |

